# IAV-CNN: a 2D convolutional neural network model to predict antigenic variants of influenza A virus

**DOI:** 10.1101/2020.07.15.204883

**Authors:** Rui Yin, Nyi Nyi Thwin, Pei Zhuang, Yu Zhang, Zhuoyi Lin, Chee Keong Kwoh

**Affiliations:** School of Computer Science and Engineering, Nanyang Technological University, Singapore, Singapore; School of Mechanical and Aerospace Engineering, Nanyang Technological University, Singapore, Singapore

## Abstract

The rapid evolution of influenza viruses constantly leads to the emergence of novel influenza strains that are capable of escaping from population immunity. The timely determination of antigenic variants is critical to vaccine design. Empirical experimental methods like hemagglutination inhibition (HI) assays are time-consuming and labor-intensive, requiring live viruses. Recently, many computational models have been developed to predict the antigenic variants without considerations of explicitly modeling the interdependencies between the channels of feature maps. Moreover, the influenza sequences consisting of similar distribution of residues will have high degrees of similarity and will affect the prediction outcome. Consequently, it is challenging but vital to determine the importance of different residue sites and enhance the predictive performance of influenza antigenicity. We have proposed a 2D convolutional neural network (CNN) model to infer influenza antigenic variants (IAV-CNN). Specifically, we introduce a new distributed representation of amino acids, named ProtVec that can be applied to a variety of downstream proteomic machine learning tasks. After splittings and embeddings of influenza strains, a 2D squeeze-and-excitation CNN architecture is constructed that enables networks to focus on informative residue features by fusing both spatial and channel-wise information with local receptive fields at each layer. Experimental results on three influenza datasets show IAV-CNN achieves state-of-the-art performance combing the new distributed representation with our proposed architecture. It outperforms both traditional machine algorithms with the same feature representations and the majority of existing models in the independent test data. Therefore we believe that our model can be served as a reliable and robust tool for the prediction of antigenic variants.

## Introduction

Seasonal influenza seriously threats public health and the global economy, causing up to 500,000 deaths and millions of cases of illness worldwide annually [1]. H1N1 and H3N2 are the principal subtypes of influenza A viruses circulating in humans [2] [3].

Vaccination is the most effective way to prevent infection and severe outcomes caused by influenza viruses [4]. The component of vaccines has to be updated regularly to ensure its efficacy [5]. The influenza virus surface glycoproteins hemagglutinin (HA) is the main target for host immunity [6]. However, the accumulation of mutations on HA proteins results in the emergence of novel antigenic variants that can not be effectively inhibited by antibodies, posing great challenges for vaccine design [7]. Developing rapid and robust methods to determine influenza antigenicity is critical to influenza vaccine design and flu surveillance.

Hemagglutinin inhibition (HI) assay is the primary method to evaluate the antigenicity of influenza viruses by measuring the ability of antisera to block the HA of the antigen from agglutinating red blood cells [8]. Smith *et al*. created an antigenic map using HI assay data and determined the antigenic evolution of influenza A H3N2 virus from 1968 to 2003 [9]. Li *et al*. developed PREDAC-H1 that systematically depicted the antigenic patterns and evolution of human influenza A H1N1 viruses [10]. By utilizing 1572 HA sequences and 197 pairs of HA sequences with HI assays data, Huang *et al*. presented the entropy and likelihood ratio to model amino acid diversity and antigenic variant score [11]. Ren *et al*. employed random forest regression and support vector regression to identify antigenicity-associated sites on HA of A/H1N1 seasonal influenza virus [12]. Richard Neher *et al*. showed a web-based application to interpret measured antigenic data and predict the properties of viruses [13]. Harvery *et al*. analyzed the sequence and 3-D structure information of HA, together with corresponding HI assay data to identify the high- and low-impact amino acid substitutions that drive the antigenic drift of influenza H1N1 viruses [14].

Numerous studies have been conducted to timely predict the antigenic variants or antigenicity of influenza viruses. Lee and Chen investigated 70 mouse monoclonal antibody binding sites for predicting antigenic variants of influenza A/H3N2 with 83% agreement [15]. Sun *et al*. provided a novel method for quantifying antigenic distance and identifying antigenic variants using sequence alone [16]. Additionally, Yin *et al*. presented a stacking model to predict antigenic variants of the H1N1 influenza virus based on epidemics and pandemics [17]. A universal computational model was integrated to predict the antigenic variants for all HA subtypes of influenza A viruses through conserved antigenic structures [18]. Regarding the prediction models on antigenicity, there are several different works to infer the influenza antigenicity with computational models. Qiu *et al*. built the antigenicity prediction model for influenza A/H3N2 viruses by incorporating the structural context of HA protein [19]. Moreover, Yao *et al*. applied a joint random forest method to human H3N2 seasonal influenza data for predicting antigenicity [20]. Zhou *et al*. presented a context-free encoding scheme of protein sequences for predicting antigenicity of diverse influenza A viruses, which encoded a protein sequence dataset into a numeric matrix and then fed the matrix into a downstream machine learning model [21]. Wang *et al*. developed a novel low-rank matrix complete model to infer antigenic distances between antigens and antisera [22]. This model exploited the correlations of the viruses and vaccines in serological tests in predicting influenza antigenicity.

Recently, deep neural networks have been successfully applied in a variety of areas including bioinformatics. Convolutional neural network (CNN) is one of the most popular approaches applied to solve bioinformatics problems, for example, classification of efflux proteins from membrane [23], human leukocyte antigen class I-peptide binding prediction [24], prediction of protein secondary structure [25] and prediction of protein-protein interaction [26]. In this paper, we leverage deep learning techniques from the natural language processing (NLP) domain to tackle the problem of antigenic variants prediction of influenza A viruses. Specifically, a new distributed representation amino acids, named ProtVec, is introduced that maps a 3-grams (three consecutive amino acids) to a 100-dimensional vector space. We then propose an approach that combines the 2D CNN model with squeeze-and-excitation mechanisms, named IAV-CNN, for the task of antigenic variants prediction. Fig. 1 illustrates the flowchart of our proposed model. The main contributions of this work are:

**Figure 1.**
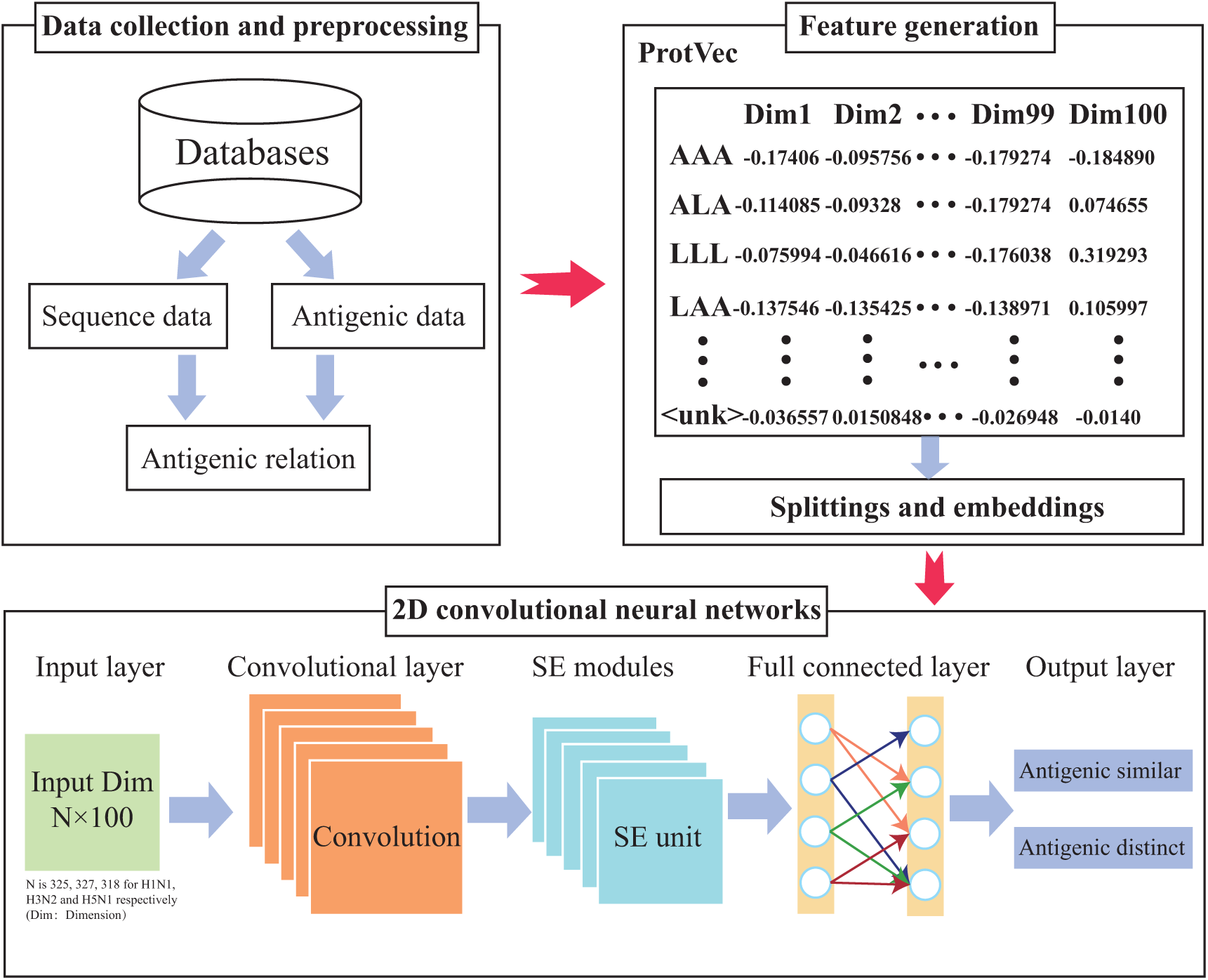
The workflow of our proposed model for predicting influenza antigenic variants using two-dimensional convolutional neural networks with squeeze-and-excitation modules.

- We propose a 2D convolutional neural network that leverages a new distributed representation of amino acids for the prediction of antigenic variants of influenza A viruses. The combination of squeeze-and-excitation units enables our models to focus on informative residues features and improve the performance.
- Extensive experiments are conducted on three public influenza datasets to evaluate the proposed model in comparison with the existing computational approaches.
- To the best of our knowledge, we perform the first attempt to predict influenza antigenicity using CNN models with new distributed representation. We believe it provides novel insights into the prediction of influenza antigenicity.

## Materials and Methods

### Dataset

In the experiment, we adopt antigenic data and sequence data of influenza subtypes H1N1, H3N2 and H5N1. The antigenic data obtained by hemagglutination inhibition (HI) assay is collected from reports of international organizations and published papers including World Health Organization (WHO), European Centre for Disease Prevention and Control (ECDC), The Francis Crick Institute (FCI), Food and Drug Administration (FDA). In total, 1562, 1249 and 666 distinct pairs of antigenic data are collected for influenza A/H1N1, A/H3N2, and A/H5N1, respectively. Correspondingly, the protein sequences of HA are derived from Influenza Virus Resource (IVR) [27] and Global Initiative on Sharing All Influenza Data (GISAID) [28]. (The information of sequences from GISAID can be found in the supplementary materials) The sequences are selected by full-length strains with the human host and duplicate sequences are eliminated from the collection. Finally, we end up with 294, 697 and 260 unique HA sequences for subtypes H1N1, H3N2 and H5N1.

### Preprocessing

The antigenic distance *D*_*ij*_ between two strains is defined by Archetti-Horsfall distance [29] as follows:

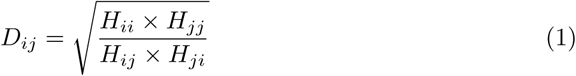

where the HI titer *H*_*ij*_ is the maximum dilution of antisera raised in strain *i* to inhibit cell agglutination caused by strain *j*. If the antigenic distance *D*_*ij*_ is equal or greater than 4, a threshold defined by Liao et al. [30], strain *i* and strain *j* are antigenic distinct. Otherwise, the pair of strains are regarded as antigenic similar. For the repetitive strain pairs where the HI titer is measured in multiple independent institutions, we utilize the median titer value to calculate the antigenic distance [31]. As a result, 937, 606, 409 antigenic distinct pairs and 625, 643, 257 antigenic similar pairs of A/H1N1, A/H3N2 and A/H5N1 strains are acquired.

For HA sequence data, we only keep the HA1 proteins for each subtype and the signal peptide is removed from the collected HA1 sequences. As a result, we obtain the HA1 sequences with the lengths of 327, 329 and 320 for H1N1, H3N2, and H5N1, respectively. Multiple sequence alignment is performed using the software MAFFT [32] on HA1 proteins for each subtype. Furthermore, the laboratory-generated reassortment sequences and the sequences with a gap ratio greater than 10% are also eliminated by a manual check. We finally obtain 294 unique sequences for H1N1, 697 for H3N2 and 260 for H5N1 in this study. The amino acid numbering of these protein sequences across different subtypes is recommended by Burke and Smith [33].

### Feature generation

The representation of biological sequences is one of the most important problems expressing the biological information with a discrete model or a vector that keeps key pattern characteristics. This is because all the existing machine learning models are only applicable to numerical vectors but not sequences as elucidated in a comprehensive review [34]. Distributed representation has displayed significant success in NLP to train word embeddings, the mapping of words to numerical vector space [35, 36]. Recently, it has been explored for bioinformatics applications such as protein classification [37] and structure prediction [38]. To convert the protein sequence information into feature sets that can be managed by neural networks, ProtVec is introduced to encode proteins through distributed representation that each trigram (sequence of three amino acids) protein is embedded in the size of 100-dimension vector [37].

To preserve the sequence pattern information, we break protein sequences into shifted overlapping residues in the window size of 3 (3-grams). The splittings and embeddings are shown in Fig 2. Here we take subtype H1N1 as an example to describe the process that a pair of influenza HA1 proteins are represented by 325 pairs of trigrams. The subtraction of a pair of trigrams characterizes the distinction between two strains at certain positions that can be denoted by a difference vector. The difference vectors *V* = [*v*_1_, *v*_2_, …, *v*_325_] are derived from ProtVec embeddings. For each vector, i.e. *v*_1_ = *ProtV ec*(*trigram*1) − *ProtV ec*(*trigram*2), where *ProtV ec*(*trigram*) is the distributed representation of a trigram in 100-dimension vector space, mapping from ProtVec. Therefore, the antigenic relationship between two HA1 strains is represented in a 325 * 100 dimensional vector space. The trigram that contains ‘-’ at any positions will be assigned the ‘unknown’ embedding from ProtVec. By formulating sequence data into distributed numerical vectors, standard machine learning algorithms can be readily applied.

**Figure 2.**
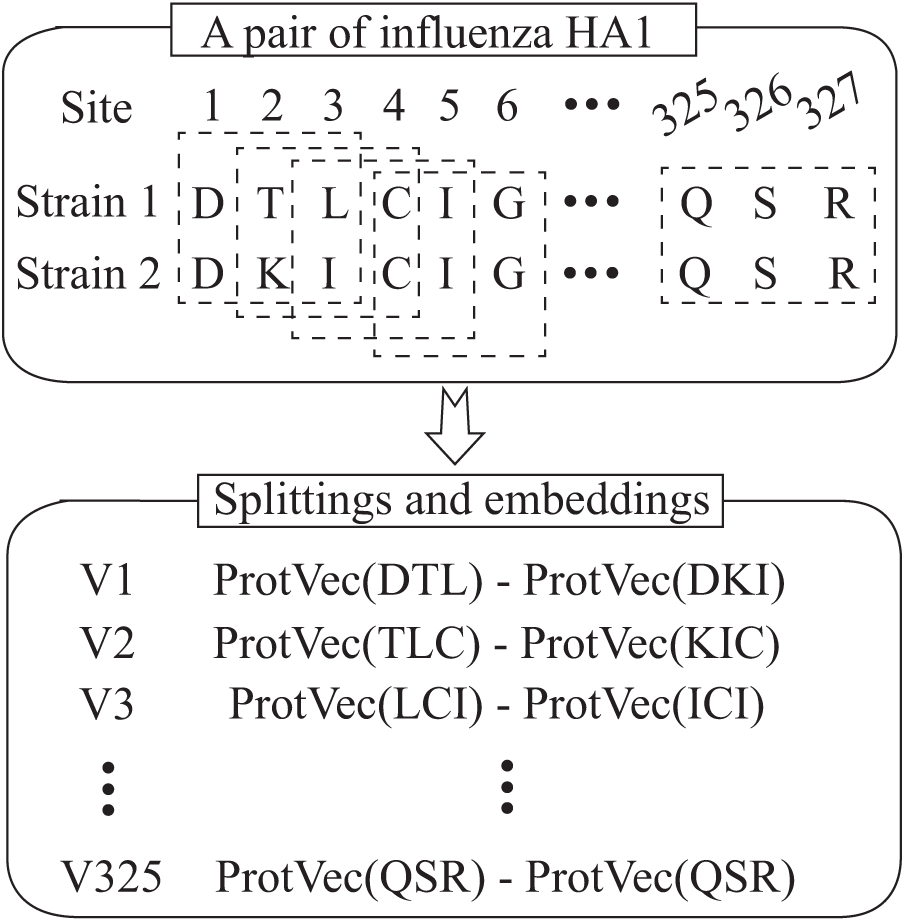
The procedure of splittings and embeddings of a pair of influenza H1N1 HA1 proteins. Each pair is embedded in a 325 * 100 dimensional vector space to represent the information of antigenic distance. Strain 1: A/California/07/2009, Strain 2: A/Ohio/9/2015.

### CNN structure

Convolutional neural networks have been applied in many fields with impressive results, especially in computer vision when the input is generally a 2D image. Much of the recent fervor has been spurred by both accessibilities to large training datasets and advances in cheap computing power to train deep neural networks in an affordable amount of time. Although originally proposed for image classification [39] [40] [41], CNN have been found work well for biological sequence data such as protein classification [42] [23] [43] and prediction of protein function [44] [45]. Encouraged by the successful application of CNN, we take advantage of the CNN architecture applied to 2D image classification and conveniently generate similar 2D inputs of the ProtVec-based matrix that explores the antigenic relationship between two influenza strains. It is with this insight that we propose IAV-CNN that aims at the task of predicting antigenic variants of influenza A virus with convolutional neural networks.

Regarding the way we construct IAV-CNN, we first follow the fundamental CNN architecture. To enhance the representational power of the network and boost meaningful sites of strains, while suppressing weak ones, the Squeeze-and-Excitation (SE) block [46] is introduced in the CNN architecture. The SE block squeezes along the spatial domain and reweights along the channels. The attention and gating mechanisms are activated by modeling the interdependencies between the channels of feature maps. The main idea is to add parameters to each channel of a convolutional block so that the network can adaptively adjust the weighting of each feature map and emphasize useful channels. Hence, we are capable of biasing the allocation of available computational resources towards the most informative residues of strains through SE blocks. The illustration of the SE block is shown in Fig 3.

**Figure 3.**
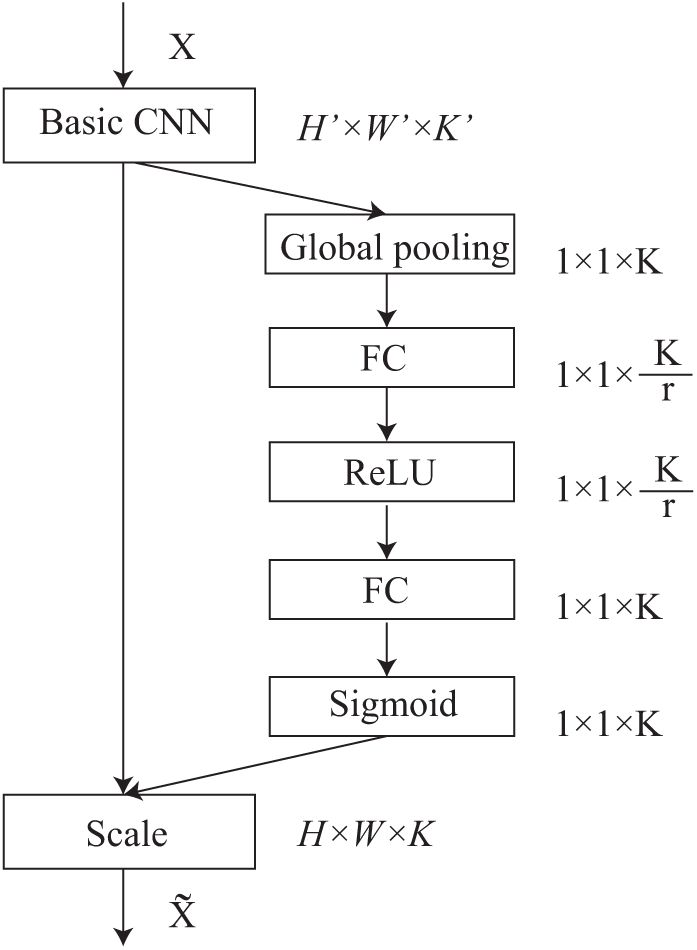
The schematic overview of squeeze-and-excitation unit with fundamental CNN module.

We assume an input *X* ∈ ℝ*H′* ×*W ‘* ×*K′* that passes through a transformation *F*_*tr*_, a convolutional operator, to generate output feature map *U* ∈ ℝ*H*×*W* ×*K*. Here *H′* and *W, H* and *W′* are the spatial height and width before and after transformation, with *K′* and *K* being the input and output channels. The vector *V* = [*v*_1_, *v*_2_, …, *v*_*K*_] represents the learned set of filter kernels, where *v*_*k*_ stands for the parameters of the *k*-th filter.

The output is denoted as *U* = [*u*_1_, *u*_2_, …, *u*_*k*_]. For each *u*_*k*_, it is formulated by

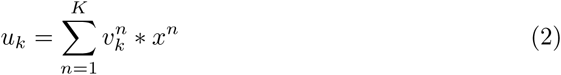

where * denotes convolution and 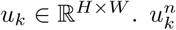 is a 2D spatial kernel denoting a single channel of *v*_*k*_ that acts on the corresponding channel of input *X*. To tackle the issue of exploiting channel dependencies, we squeeze global spatial information into a channel descriptor by using global average pooling to generate channel-wise statistics. Consequently, a statistic *z* is obtained by squeezing *U* through its spatial dimensions *H* × *W*. The *k*-th element of *z* is formulated by

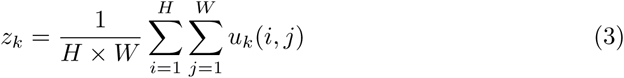

To capture channel-wise dependencies and make full use of information aggregated in the squeeze operation, we employ a gating mechanism with a sigmoid activation. The equation is described below, where *δ* refers to the ReLU function [47], 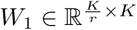 and 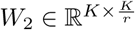. The gate mechanism consists of two full-connected (FC) layers around the non-linearity, i.e. a ReLU and then follow by sigmoid activation, which returns to the channel dimension of the transformation output *U*. The hyperparameter *r* is the reduction ratio that allows us to adjust the computational cost and capacity of the SE modules in the network [46].

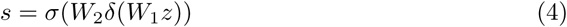

The output of the SE module is finally obtained by rescaling *U* with the activation *s*

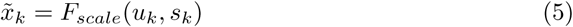

Where 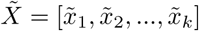 and *F*_*scale*_(*u*_*k*_, *s*_*k*_) is the multiplication between the scalar *s*_*k*_ and the feature map *u*_*k*_. In this regard, the squeeze operator compresses global spatial information into local descriptors and the excitation operator maps these specific descriptors into a set of channel weights. Consequently, SE modules present a global understanding of each channel by squeezing the feature maps to a single numeric value and dynamically change it by adding a content-aware mechanism to weight each channel. Algorithm 1 clarifies the detailed steps of our proposed model for predicting influenza antigenic relationships using 2D convolutional neural networks based on ProtVec.

#### Algorithm 1 The IAV-CNN algorithm for predicting influenza antigenic variants through ProtVec.

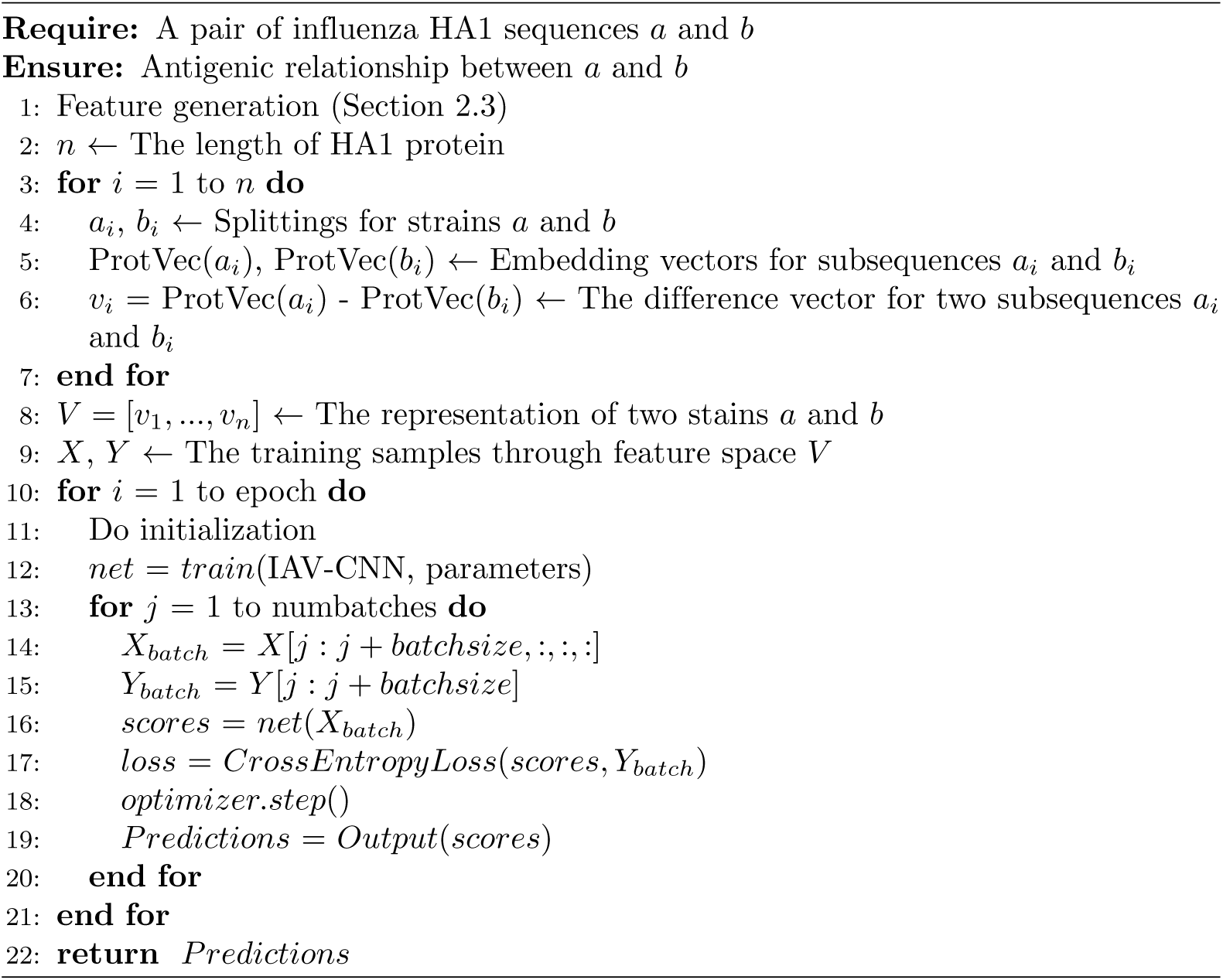

### Experiments

#### Baseline Approaches

We set up two types of baselines to evaluate our model. The first baseline is we compare our proposed model with several traditional machine learning algorithms using the same feature space for the prediction tasks. The classifiers include logistic regression (LR), K-nearest neighbor (KNN), support vector machine (SVM), random forest (RF), neural network (NN) and CNN model without SE blocks. The second baseline is to apply several art-of-the-state approaches. Liao et al. proposed a method by incorporating scoring and regression methods to predict antigenic variants [30]. Yao et al. developed a joint random forest regression algorithm, cooperatively consider top substitution matrices that can improve the prediction performance [20]. Lee and Chen used the number of amino acid changes located on the five epitope regions for the antigenic variants prediction [15]. Lees et al. provided an update for the frequently referenced five antigenic sites and increase additional assignments to establish five canonical regions [48]. They constructed a range of linear models based on banded changes for the prediction. Peng et al. built a universal model for the antigenic variation prediction of influenza A virus H1N1, H3N2 and H5N1 using conserved antigenic structures [18]. We will reconstruct these models to predict antigenic variants on three influenza datasets in comparison with our proposed algorithm.

#### Implementation and evaluation

All the approaches are implemented with Scikit-learn [49] and PyTorch [50]. The antigenic distinct is labeled as ‘1’ and antigenic similar is ‘0’ for the relationship of two strains. The influenza datasets of each subtype are randomly divided into independent training and testing set with a ratio of 0.8:0.2. We construct our model and evaluate the training process on the training dataset and the independent testing dataset is used to assess its capability in predicting relations of novel antigenic variants. For CNN-based models, we apply several algorithms with a minimum batch size of 32 for optimization. The one that achieves the best performance of the experimental results will be selected. The drop-out (rate=0.5) strategy is carried out with the learning rate of 0.001 and all the models are iterated for 100 training epochs. We adopt five different metrics including accuracy, precision, recall, f-score and Matthews’s Correlation Coefficient (MCC) to evaluate the predictive performance of the models. Accuracy describes the proportion of true results among the total number of samples. Precision reveals the proportion of predicted positives that are truly positive. Recall indicates the proportion of actual positives correctly classified. The f-score is the harmonic mean of precision and recall that maintains a balance between the two metrics [51]. MCC is used as a measure of the quality of classifications in machine learning that is less influenced by imbalanced test sets since it considers mutually accuracies and error rates on both classes [52].

## Results and discussion

The quality and reproducibility of the model is a crucial factor for the study. In the experiments, we first investigated the effect of using different optimizers including Adaptive Moment Estimation (Adam) [53], Adadelta [54], Adaptive Gradient Algorithm (AdaGrad) [55], Root Mean Square Propagation (RMSProp) [56] and Stochastic Gradient Descent (SGD) [57] on our model. Next, we described our model and how it exerted a new distributed representation of amino acids to solve the problem of antigenicity prediction over other traditional classifiers. Finally, the comparative performance between IAV-CNN and several recently developed state-of-the-art methods is presented to further validate the ability of our model.

### The performance of IAV-CNN with different optimizers

Table 1 shows the predictive performance of IAV-CNN with five different optimizers on the testing data of three influenza subtypes. The best results for each dataset are highlighted in bold. We can observe from the table that by using SGD optimizer, it achieves the best performance of 0.856, 0.873, 0.861 0.867 and 0.656 in terms of accuracy, precision, recall, F-score and MCC on H3N2 influenza data. Similarly, when applied SGD optimizer in the other two datasets, our model also displays the best performance in all of the metrics except recall. Therefore, we use SGD algorithm as the optimizer on subsequent experiments in comparison with other approaches for antigenicity prediction. However, varied performance is observed for different datasets, for instance, H1N1 displays an overall more desirable outcome than the other two types, This may largely owe to the inconsistent sample size that the model on H1N1 dataset presents better predictive results compared with H3N2 and H5N1.

**Table 1.** Performance comparison of IAV-CNN model with different optimizers on H1N1, H3N2 and H5N1 datasets.

| Dataset | Optimizer | Accuracy | Precision | Recall | F-score | MCC |
| --- | --- | --- | --- | --- | --- | --- |
| H1N1 | Adam | 0.850 | 0.857 | 0.914 | 0.885 | 0.663 |
|  | Adadelta | 0.856 | 0.928 | 0.824 | 0.873 | 0.716 |
|  | AdaGrad | 0.885 | 0.896 | 0.915 | 0.906 | 0.759 |
|  | RMSProp | 0.872 | 0.871 | **0.933** | 0.901 | 0.725 |
|  | SGD | **0.917** | **0.928** | 0.915 | **0.924** | **0.806** |
| H3N2 | Adam | 0.796 | 0.866 | 0.829 | 0.792 | 0.603 |
|  | Adadelta | 0.806 | 0.831 | 0.837 | 0.847 | 0.601 |
|  | AdaGrad | 0.828 | 0.851 | 0.859 | 0.855 | 0.622 |
|  | RMSProp | 0.792 | 0.846 | 0.824 | 0.793 | 0.598 |
|  | SGD | **0.856** | **0.873** | **0.861** | **0.867** | **0.656** |
| H5N1 | Adam | 0.836 | 0.868 | 0.846 | 0.857 | 0.665 |
|  | Adadelta | 0.836 | 0.818 | 0.878 | 0.867 | 0.662 |
|  | AdaGrad | 0.843 | 0.880 | 0.846 | 0.863 | 0.681 |
|  | RMSProp | 0.851 | 0.882 | **0.889** | 0.870 | 0.701 |
|  | SGD | **0.881** | **0.908** | 0.885 | **0.896** | **0.756** |

### Comparative performance between IAV-CNN and traditional classifiers on ProtVec-based features

We further examine the performance using several classical algorithms for predicting antigenic variants with ProtVec-based features on three influenza datasets. Table 2 summarizes the comparative results of IAV-CNN and other traditional machine learning methods including logistic regression, k-nearest neighbor, support vector machine, random forest and neural network. For a fair comparison, we use the optimal parameters for the classifiers in all experiments. Specifically, for all subtypes of influenza data, it is observed that random forest and neural networks perform higher accuracy than our proposed model on the training data, whereas IAV-CNN has demonstrated better predictive results on the testing data. This is probably the overfitting problem that the classic algorithms fit too well with the training data. It then becomes difficult for the models to generalize to new samples that are not in the training set. Our proposal model overcomes this issue by applying the dropout mechanism that randomly sets activations to zero during the training process to avoid overfitting. We obtain the accuracy of 0.917, 0.856 and 0.881 for three subtypes. The results are 5.4%, 6.4% and 5.3% higher than the best traditional classifiers, which only achieve 0.863, 0.792 and 0.828, respectively. Besides, we have noticed that the simple SVM algorithm is not suitable for the prediction of small-scale H5N1 data. The SVM finds a maximum edge hyperplane for classification. Since there is no large number of the iterative process, the prediction ability is limited and the accuracy is low.

**Table 2.** Comparative performance between IAV-CNN and other machine learning methods using ProtVec features on training and testing data of three influenza subtypes.

| Subtype | Model | Training data |  |  |  |  | Testing data |  |  |  |  |
| --- | --- | --- | --- | --- | --- | --- | --- | --- | --- | --- | --- |
|  |  | Accuracy | Precision | Recall | F-score | MCC | Accuracy | Precision | Recall | F-score | MCC |
| H1N1 | LR | 0.817 | 0.816 | 0.892 | 0.853 | 0.616 | 0.722 | 0.752 | 0.826 | 0.787 | 0.392 |
|  | KNN | 0.901 | 0.956 | 0.873 | 0.913 | 0.803 | 0.815 | 0.915 | 0.774 | 0.839 | 0.637 |
|  | SVM | 0.594 | 0.594 | 1.000 | 0.745 | 0.409 | 0.623 | 0.623 | 1.000 | 0.768 | 0.423 |
|  | RF | 0.987 | 0.993 | 0.985 | 0.989 | 0.974 | 0.863 | 0.884 | 0.897 | 0.891 | 0.706 |
|  | NN | 0.998 | 0.997 | 0.999 | 0.998 | 0.995 | 0.859 | 0.895 | 0.877 | 0.886 | 0.703 |
|  | CNN | 0.978 | 0.986 | 0.970 | 0.978 | 0.952 | 0.880 | 0.896 | 0.928 | 0.912 | 0.761 |
|  | IAV-CNN | 0.968 | 0.976 | 0.972 | 0.974 | 0.937 | 0.917 | 0.928 | 0.915 | 0.924 | 0.806 |
| H3N2 | LR | 0.847 | 0.872 | 0.793 | 0.831 | 0.694 | 0.696 | 0.761 | 0.624 | 0.686 | 0.404 |
|  | KNN | 0.863 | 0.893 | 0.808 | 0.848 | 0.727 | 0.728 | 0.804 | 0.647 | 0.717 | 0.471 |
|  | SVM | 0.473 | 0.473 | 1.000 | 0.643 | 0.403 | 0.532 | 0.532 | 1.000 | 0.695 | 0.429 |
|  | RF | 0.962 | 0.968 | 0.951 | 0.959 | 0.924 | 0.776 | 0.824 | 0.737 | 0.778 | 0.557 |
|  | NN | 0.973 | 0.967 | 0.977 | 0.972 | 0.946 | 0.792 | 0.817 | 0.737 | 0.775 | 0.548 |
|  | CNN | 0.961 | 0.975 | 0.972 | 0.973 | 0.962 | 0.841 | 0.866 | 0.854 | 0.860 | 0.621 |
|  | IAV-CNN | 0.968 | 0.975 | 0.973 | 0.974 | 0.950 | 0.856 | 0.873 | 0.861 | 0.867 | 0.656 |
| H5N1 | LR | 0.889 | 0.902 | 0.921 | 0.912 | 0.763 | 0.813 | 0.863 | 0.808 | 0.834 | 0.623 |
|  | KNN | 0.883 | 0.930 | 0.879 | 0.904 | 0.758 | 0.799 | 0.849 | 0.795 | 0.821 | 0.593 |
|  | SVM | 0.378 | 0.000 | 0.000 | 0.000 | 0.000 | 0.418 | 0.000 | 0.000 | 0.000 | 0.000 |
|  | RF | 0.976 | 0.991 | 0.970 | 0.980 | 0.949 | 0.828 | 0.867 | 0.833 | 0.850 | 0.651 |
|  | NN | 0.981 | 0.997 | 0.973 | 0.985 | 0.961 | 0.828 | 0.867 | 0.833 | 0.850 | 0.651 |
|  | CNN | 0.978 | 0.995 | 0.992 | 0.973 | 0.962 | 0.841 | 0.866 | 0.854 | 0.860 | 0.621 |
|  | IAV-CNN | 0.955 | 0.997 | 0.990 | 0.993 | 0.977 | 0.861 | 0.882 | 0.870 | 0.876 | 0.715 |

### Comparative performance between IAV-CNN and other methods

To demonstrate the effectiveness of our model, we compared IAV-CNN with several state-of-the-art methods on the prediction of influenza antigenicity on three datasets. Cross-validation is often leveraged to examine a predictor for its capability in practical applications. Here we adopt the 5-fold cross-validation in the training data that has been utilized by many investigators to construct the predictive models and evaluate our model on the remaining testing data. According to the experimental results in Table 1 and 2, SGD has been chosen as the best optimizer for our model. We still use the default learning rate (0.001) with a dropout (0.5) mechanism in the experiments for CNN based models. Furthermore, independent testing data is used to evaluate the ability of our model to predict new sample data with robustness. Fig 4 shows the performance comparison of IAV-CNN with other state-of-the-art methods on independent testing data, as detailed below.

**Figure 4.**
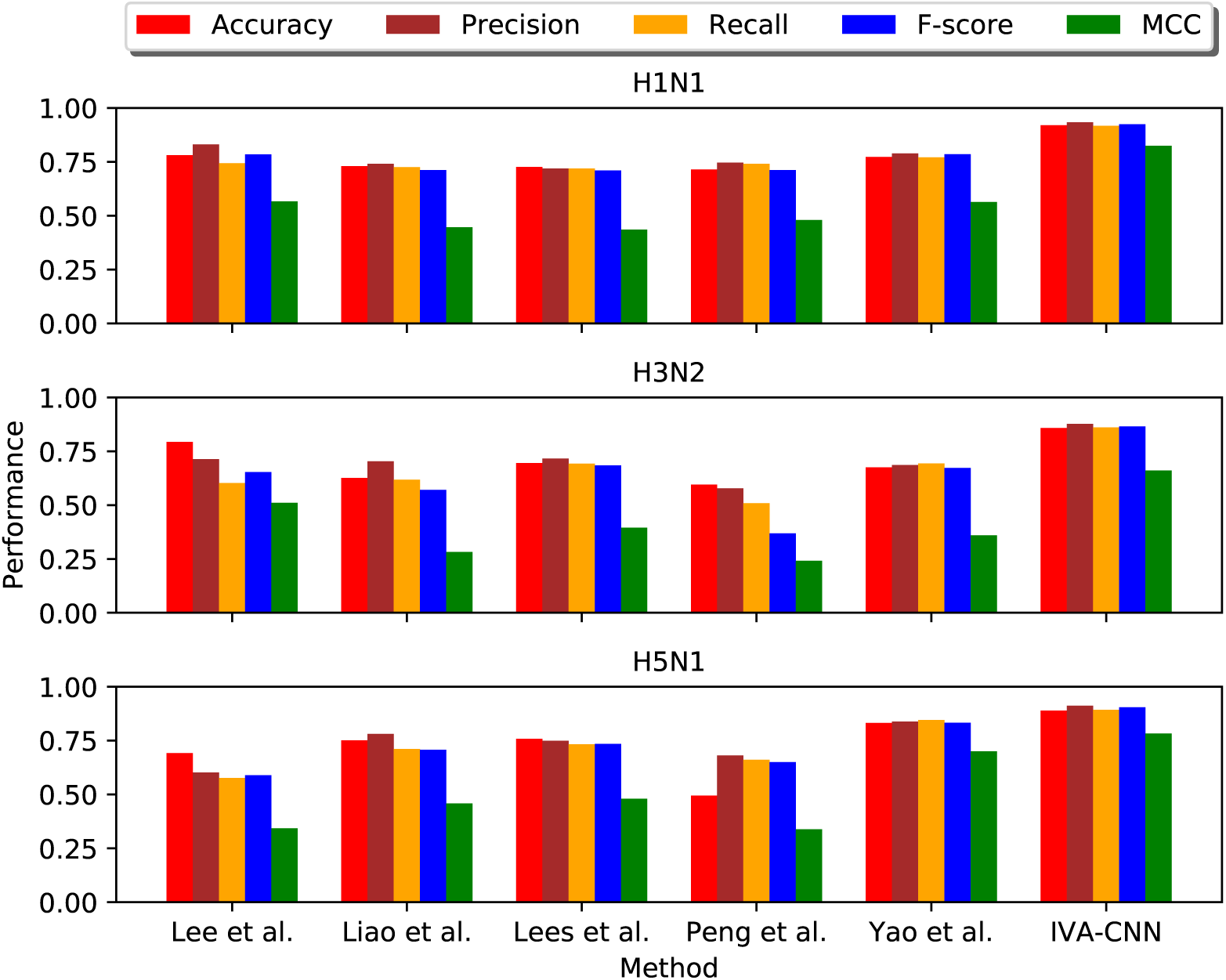
The comparative performance between IAV-CNN and other state-of-the-art methods for predicting influenza antigenic variants on independent testing data of three influenza subtypes.

The x-axis represents the different methods we applied for the prediction and the y-axis shows the values of all metrics. It is shown that our proposed model achieves a remarkable higher performance than compared methods. In more detail, IAV-CNN can obtain an accuracy of 0.920, 0.858 and 0.889 for independent H1N1, H3N2 and H5N1 testing data, respectively. The results are 13.9%, 6.5% and 6.7% higher than the best performance of compared methods. Regarding other evaluation metrics, the results also indicate that IAV-CNN outperforms the current state-of-the-art methods on all datasets. Overall, it is demonstrated that IAV-CNN can accurately predict influenza antigenic variants on selected subtypes with feasibility and robustness. It may also be applicable to predict the antigenicity of a wide range of viruses and drive the development of personalized medicine for infectious diseases.

### Interpretation

The prediction of influenza antigenicity is critical for the study of viral evolution and vaccine selection. Although many methods have been proposed to predict novel influenza variants using diverse feature representations, i.g. epitope and physicochemical properties, when establishing the machine learning models, the correlation between features are never taken into consideration. Our proposed IAV-CNN is an important type of 2D CNN model consisting of a convolutional kernel, squeeze-and-excitation module and a full-connected layer. By utilizing IAV-CNN, we try to capture meaningful residue sites and even hidden features by scanning the sequences of pair of strains. The introduction of SE modules helps us to focus on the sites with different residues that are given larger weights in the training process. The results prove that IAV-CNN can enhance the predictive performance over other existing machine learning approaches by capturing important residue sites of the compared strains. As a result, our proposed model can be served as a reliable tool for the prediction of influenza antigenicity, which assists biologists to gain a better understanding of influenza evolution and vaccine selection.

## Conclusion

In this paper, we have described the feasibility of applying machine learning techniques from NLP domain to solve bioinformatics problems, particularly, the antigenicity prediction of influenza A viruses. We propose IAV-CNN to extract a vector space with a distributed representation of amino acids through ProtVec and predict the influenza antigenic variants, using a 2D CNN architecture with squeeze-and-excitation mechanisms. Compared with other traditional machine learning algorithms, IAV-CNN produces superior predictive efficacy with the same feature representations on three different influenza datasets. Moreover, further experiments demonstrate our model achieves state-of-the-art antigenicity prediction results on the majority of test sets over existing models. We believe this framework is capable of making predictions for any subtypes of influenza viruses with sufficient training data, and facilitate flu surveillance.

## Supporting information

The codes and data to generate the IAV-CNN are publicly available at: https://github.com/Rayin-saber/IAV-CNN

## Acknowledgments

This project is supported by AcRF Tier 2 grant MOE2014-T2-2-023, Ministry of Education, Singapore and A*STAR-NTU-SUTD AI Partnership grant, RGANS1905.

